# Structural and Functional Brain-wide Alterations in A350V IQSEC2 Mutant Mice Displaying Autistic-like Behavior

**DOI:** 10.1101/2020.09.05.284364

**Authors:** Daniela Lichtman, Eyal Bergmann, Alexandra Kavushansky, Nadav Cohen, Nina S. Levy, Andrew P. Levy, Itamar Kahn

## Abstract

IQSEC2 is an X-linked gene which is associated with autism spectrum disorder (ASD), intellectual disability and epilepsy. IQSEC2 is a postsynaptic density protein, localized on excitatory synapses as part of the NMDA receptor complex and is suggested to play a role in AMPA receptor trafficking and mediation of long-term depression. Here, we present brain-wide structural volumetric and functional connectivity characterization in a novel mouse model with a missense mutation in the IQ domain of IQSEC2 (A350V). Using high-resolution structural and functional MRI, we show that animals with the A350V mutation display increased whole-brain volume which was further found to be specific to the cortex and hippocampus. Moreover, using a data-driven approach we demonstrate that A350V mice present alterations in structure–function relations of the frontal, auditory, and visual networks. Examination of these alterations revealed an increase in functional connectivity between the anterior cingulate cortex and the dorsomedial striatum. We also show that corticostriatal functional connectivity is correlated with individual variability in social behavior only in A350V mice, as assessed using the three–chamber social preference test. Our results at the systems-level bridge the impact of previously reported changes in AMPA receptor trafficking to network-level disruption and impaired social behavior. Further, the A350V mouse model recapitulates similarly reported brain-wide changes in other ASD mouse models, with substantially different cellular-level pathologies that nonetheless result in similar brain-wide alterations, suggesting that novel therapeutic approaches in ASD that result in systems-level rescue will be relevant to IQSEC2 mutations.

**Significance Statement:** Several recent studies have characterized the changes in the organization of brain networks in animal models of autism spectrum disorders (ASD). Here we assessed the effect of an A350V missense mutation in the IQSEC2 gene, which is associated with ASD, on brain-wide functional connectivity and its relation to social behavior deficits in A350V mice relative to controls. We found that the A350V IQSEC2 model results in disrupted functional connectivity of the anterior cingulate cortex and the dorsomedial striatum. Critically, disrupted increased corticostriatal functional connectivity is predictive of individual variability in social interaction only in A350V mice implicating this pathway in the pathophysiology of the A350V IQSEC2 mutation.

## Introduction

Autism spectrum disorder (ASD) is a neurodevelopmental disorder with a highly genetically heterogeneous component (Geschwind and State, 2015; Takata et al., 2018). It is diagnosed on the basis of a combination of behavioral observations and clinical interviews that assess deficits in social interactions, communication and language, as well as repetitive and stereotyped behaviors (Elsabbagh et al., 2012), and is found to be associated with other cognitive and neurological conditions (Matson and Nebel-Schwalm, 2007; Mannion and Leader, 2013). Despite the complexity of ASD, genetic research has contributed significantly to the elucidation of its pathophysiology (Geschwind and State, 2015). Research in genetically heterogeneous ASD populations has demonstrated converging pathophysiology, pointing out the role of glutamatergic cortical synapses in the pathology (Gilman et al., 2011; Parikshak et al., 2013). At the systems level, findings in human and genetic animal models have implicated disrupted brain-wide connectivity in ASD pathophysiology with diametrically different effects on distal versus local connectivity (Liska and Gozzi, 2016; Vasa et al., 2016).

Intrinsic functional connectivity MRI (fcMRI), the temporal correlation of spontaneous blood oxygenation level-dependent (BOLD) signal fluctuations in the brain, has been shown to be a useful method for characterization of brain networks in humans and animals (Fox and Raichle, 2007; Power et al., 2014b; Stafford et al., 2014; Zerbi et al., 2015; Bergmann et al., 2016). Several studies have demonstrated that fcMRI can be used to identify functional alterations in human diseases and in transgenic models in rodents (Buckner et al., 2009; Liska and Gozzi, 2016; Bertero et al., 2018; Shofty et al., 2019). Specifically, fcMRI studies in several ASD animal models have detected functional connectivity alterations that are consistent with electrophysiological recordings and human fcMRI studies (Scott-Van Zeeland et al., 2010; Peixoto et al., 2016; Liska et al., 2018; Pagani et al., 2019; Shofty et al., 2019).

IQSEC2, an X-linked gene coding for a protein found in the post synaptic density of glutamatergic synapses, has been implicated in trafficking of a-amino-3-hydroxy-5-methyl-4-isoxazolepropionic acid (AMPA) receptors and regulation of synaptic transmission (Murphy et al., 2006; Brown et al., 2016; Petersen et al., 2018; Rogers et al., 2019). In humans, mutation in the IQSEC2 gene is associated with ASD, intellectual disabilities and epilepsy (Shoubridge et al., 2010; Kalscheuer et al., 2016; Zipper et al., 2017; Mignot et al., 2019). We recently described a new mouse model (Rogers et al., 2019) with a missense mutation in the IQ domain of IQSEC2 at amino acid residue 350 resulting in a valine for alanine substitution (A350V). The A350V mouse model was generated using CRISPR technology based on a human *de novo* mutation (Zipper et al., 2017) and demonstrated a significant reduction in the surface expression of GluA2 AMPA receptors in hippocampal neurons along with increased locomotion activity and abnormal social behavior, demonstrating that it manifests some of the abnormalities found in the human condition that inspired the generation of this mouse model.

Here, we investigated the impact of the A350V mutation on brain-wide structural and functional alterations using MRI. We examined the A350V mouse model using a structure–function analysis that allowed us to characterize putative changes in functional connectivity in a hypothesis-free data-driven approach. Further, we tested whether the observed functional connectivity alterations can explain behavioral variability, thereby linking functional connectivity changes to ASD-related behavioral impairments. By characterizing the discrepancies between structure–function relations and further investigating the altered regions, we found that the A350V mouse model presents increased corticostriatal functional connectivity which is linked to abnormal social behavior in the three-chamber sociability task.

## Materials and Methods

### Ethics

All animal experiments were conducted in accordance with the United States Public Health Service’s Policy on Humane Care and Use of Laboratory Animals and approved by the Institutional Animal Care and Use Committee of the Technion – Israel Institute of Technology.

### Animals and housing conditions

A350V IQSEC2 mice in a C57Bl/6J background were generated by CRISPR as previously described (Rogers et al., 2019). A350V IQSEC2 hemizygous male mice and male wild-type (WT) littermates were housed in groups of 2–5 animals per cage in a reversed 12 h light-dark cycle with food and water available *ad libitum*. The housing room was maintained at 23 ± 2 °C. All experiments were conducted during the dark phase.

### Experimental design

The experimental protocol is presented in **Fig. 1**. Behavioral experiments were performed on 6–7-week-old mice in which all animals were tested for social preference (*n*_WT_ = 18, *n*_A350V_ = 14). A subset of the cohort then underwent a head-post surgery at 10–12 weeks of age and were allowed to recover for one week in their home cage. Two mice (one of each genotype) died during the surgery and therefore two additional mice, who did not participate in the behavioral experiments, were operated on, to replace these mice (*n*_WT_ = 13, *n*_A30V_ = 13). To reduce head movement during the fcMRI scans, mice were acclimatized to the head fixation position inside the scanner for four days prior to the first fcMRI scan. Thus, at 3–4 months of age the mice underwent 7–8 awake head-fixed fcMRI sessions (one session per day) which were followed with a single high-resolution structural imaging scan under anesthesia on the last fcMRI session.

**Figure 1.**
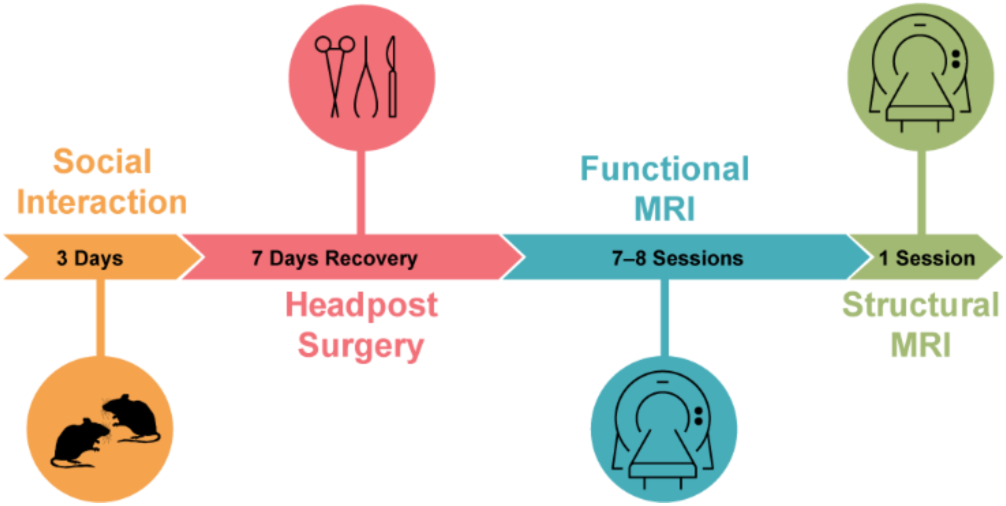
Experimental protocol. The timeline shows the sequence of experimental steps animals underwent. Social interaction assessment was examined on juvenile mice at 6–7 weeks of age. Then, at 10–12 weeks of age mice underwent a head-post surgery and recovered for seven days. After recovery, mice underwent a four days acclimatization period to awake head-fixed imaging followed by a 7–8 fcMRI sessions. Finally, a single structural MRI scan was acquired.

### Three-chamber sociability test

Social interaction was measured in order to assess autistic-like behavior using the three-chamber test (Moy et al., 2004). Subject mice were assessed for the tendency to prefer an unfamiliar conspecific mouse (social stimulus; *Stranger 1*) over a novel object. Mice (subject and stimulus) were habituated to the testing room for three consecutive days prior to the test day for one hour. Stimulus mice were further habituated to the wire cages (10.8 cm in height and 10.2 cm diameter at the bottom; Galaxy Cup, Spectrum Diversified Designs, Inc., Streetsboro, OH, USA) for 20 min each day. On the test day, subject mice were habituated to the apparatus (70 × 29 × 35 cm^3^) for 10 min and were able to explore all three empty chambers. Time spent in each chamber was measured to assess chamber bias. Following habituation, subject mice were assessed for social preference for 10 min by allowing interaction with *Stranger 1* which was placed inside a wire cage in one chamber and a novel object which was placed inside an identical wire cage in the opposite chamber. Stimulus mice location was counterbalanced across trials to prevent chamber bias. Stimulus mice were conspecific C57BL/6 mice from different litters, and were age, sex and weight-matched to the subject mice and to each other.

Behavioral experiments were performed under red lighting conditions (<5 lux) by a single experimenter blind to the genotype of the animals. All experiments were video-recorded by a camera (GUPPY PRO F-125B CCD, Allied Vision Technologies, GmbH, Ahrensburg, Germany) located above the arena and analyzed post-hoc using Ethovision XT 10.1 software (Noldus, Wageningen, The Netherlands). The apparatus was cleaned after each trial with 70% ethanol and then with double-distilled water.

### Head-post surgery

To prepare for awake fMRI scanning, mice were implanted with MRI-compatible head-posts, as previously described (Bergmann et al., 2016). Briefly, mice were anesthetized with isoflurane (1.5– 2.5%), the scalp and periseptum were removed from above the surface of the skull and a head-post was attached to the skull using dental cement (C&B Metabond, Parkell, Brentwood, NY, United States). Mice received a subcutaneous injection containing broad-spectrum antibiotics (Cefalexin sodium) and Buprenorphine daily for at least two days after the surgery

### Animal imaging and analyses

MRI scans were acquired with a 9.4T MRI (Bruker BioSpin GmbH, Ettlingen, Germany), using a quadrature 86 mm transmit-only coil and a 20 mm loop receive-only coil (Bruker). Raw data were reconstructed using ParaVision 5.1 (Bruker). For both structural and functional scans, data were registered to a downsampled version of the Allen Mouse Brain Connectivity (AMBC) atlas which includes anatomical annotations (Lein et al., 2007) and optical density maps from anatomical tracing experiments (Oh et al., 2014).

### Structural MRI acquisition and analysis

Mice underwent a single anatomical scan while being anesthetized with isoflurane (0.5–1%) using a rapid relaxation enhancement (RARE) sequence (repetition time [TR] 6000 ms, echo time [TE] 8.8 ms, FA 180°, RARE factor 16, 36 coronal slices, matrix 160 × 160, field of view [FOV] 16 × 16 mm^2^, 100 × 100 × 400 μm^3^).

To estimate the structural changes in the A350V mouse model, a Deformation-based morphometry (DBM) was used by which the structural MRI scans were registered to a target using a small-displacement non-linear registration software (Andersson et al., 2007) and the Jacobian determinants were computed to serve as a measure of deformation. The target was generated by averaging the anatomical scans of WT mice (*n* = 12). To allow accurate estimation of brain volume, structural scans were kept in native space and the AMBC atlas was aligned to the target. Whole-brain volume, excluding the olfactory bulb and cerebellum, was manually defined for each animal and a whole-brain template was generated by averaging the manual labels for WT mice only. We used AMBC anatomical annotations to define the regions: cerebral cortex excluding the olfactory bulb (CTX), hippocampal region (HIP), thalamus (TH), caudoputamen (CP) and piriform cortex (PIR) and thereby allowing an estimation of anatomical changes between the groups. We further defined six cortical modules: Prefrontal, Anterolateral, Somatomotor, Visual, Medial and Temporal based on Harris et al. (2018) with the exception of the rostrolateral visual area (VISrl) which is part of the Medial module and not the Visual module as originally defined.

### Resting-state fMRI acquisition and preprocessing

Mice of each genotype (*n* = 13 per group) underwent multiple awake head-fixed fcMRI sessions, as previously described in Bergmann et al. (2016; see also Shofty et al., 2019). For each session, blood oxygenation level-dependent (BOLD) contrast scans were acquired using spin echo-echo planar imaging (SE-EPI) sequence (TR 2500 ms, TE 18.398 ms, FA 90°, 30 coronal slices, matrix 96 × 64, FOV 14.4 × 9.6 mm^2^, 150 × 150 × 450 μm^3^). Animals were scanned for 6–8 sessions, four runs per session and 200 repetitions per run. There were no significant differences in the number of included sessions between A350V (6.461 ± 1.126 [mean ± SD]) and WT mice (6.846 ± 0.688; unpaired *t*-test; *t*(24) = 1.05, *p* = 0.304).

Raw data were preprocessed as previously described (Bergmann et al., 2016, Shofty et al., 2019), including removal of the first two volumes for T1-equilibration effects, compensation of slice dependent time shifts, rigid body correction for head motion, registration to a downsampled version of the AMBC atlas (Lein et al., 2007, Oh et al., 2014), and intensity normalization. Data scrubbing was applied as previously described (Power et al., 2014a; Bergmann et al., 2016) with exclusion criteria of 50 μm framewise displacement and derivative RMS variance over voxels (DVARS) of 150% inter-quartile range (IQR) above the 75th percentile and exclusion of one frame after the detected motion. Sessions with fewer than 60 included frames were excluded. The data subsequently underwent demeaning and detrending, nuisance regression of six motion axes, ventricular and white matter signals and their derivatives, temporal bandpass filtering to retain frequencies between 0.009–0.08 Hz and spatial smoothing with a full width at half maximum of 600 μm.

### Functional connectivity analysis

To estimate functional connectivity, a region of interest (ROI, also termed seed) was defined (described below) and its time course was extracted. Seed-based Fisher’s *Z* transformed Pearson’s *r* correlation maps (*Z*(*r*)) were averaged across sessions to provide a subject-specific map. For group-level analyses, the subject-specific maps were submitted to a one-way *t*-test for each animal group separately or a two-way *t*-test when comparing the groups to each other (SPM, Wellcome Department of Cognitive Neurology, London, UK). In addition, seed-to-seed correlations were calculated and averaged across sessions to estimate functional network alterations between two defined regions and to further examine whether functional connectivity was correlated with each animal’s behavior.

### Structure–function analysis

To estimate brain-wide alterations in the A350V mouse model in a data-driven approach the overlap between functional and anatomical connectivity was tested by comparing the optical density maps with the fcMRI maps, as elaborately described in Bergmann et al. (2016; see also Stafford et al., 2014 and Grandjean et al., 2017). This analysis approach allows to test the hypothesis that functional alterations in the mutant group will result in altered structure-function relations. An advantage of this specific analysis approach is that it is not threshold dependent. For group analysis, seed-based statistical parametric maps were used to estimate anatomical prediction of functional connectivity in association and sensory regions. Functional connectivity was derived from the A350V and WT acquired here while anatomical connectivity was derived from C57BL/6J animals who were used to construct the AMBC atlas (Oh et al., 2014). The location of the seeds (450 μm-diameter spheres) for sensory (*n* = 24) and association (*n* = 13) regions were defined at the center of the injection site. Correlation maps (*Z*(*r*)) were extracted based on seeds (a three-dimensional cross from seven voxels was generated) which corresponds to 101 out of the 122 C57BL/6J injection sites that were used to define the six cortical modules in Harris et al. (2018); missing injections include 18 out of 35 injections in VISp, one in MOp and one in MOs. To characterize structure–function relations we conducted a series of receiver operating characteristic (ROC) analyses as previously described (Bergmann et al., 2016). Briefly, anatomical projection volumes were taken from Oh et al. (2014) and compared to functional volume distributions over multiple statistical thresholds (158 when comparing between the sensory and association systems and 265 when comparing between A350V and WT). A binary volume threshold of 0.05 mm^3^ was used to define true anatomical connections to examine the sensitivity and specificity of the prediction of anatomical connections based on functional connections. To estimate the accuracy of prediction, we calculated the area under each ROC curve, which estimates the relationship between true-positive and false-positive predictions at different thresholds, indicating how well functional connectivity discriminates between anatomically connected and unconnected regions.

To further examine the functional alterations of regions with altered structure–function relations, a Sørensen–Dice similarity coefficient was calculated (*D* = 2 × (A ∩ B) / (|A| + |B|)) and used to quantify the overlap between two tracer injections within the same cortical region. Then, seed-based analysis was computed to further assess intergroup differences using unpaired two-tailed Student’s *t*-test.

### Statistical analyses

All data were analyzed using MATLAB R2018a (The Mathworks, Natick, MA, USA) except for behavioral data in which analysis of variance (ANOVA) was analyzed using jamovi Version 1.0.7 (The jamovi project, 2020). The Lilliefors test was used to determine normality. Behavioral data were found to be normally distributed. As for the MRI data, since part of the data were not normally distributed, group comparisons were performed using the non-parametric statistics Mann–Whitney U-test and Wilcoxon signed-rank test. Correspondence between manual and automated volumetric measurements and correlation between fcMRI alteration and social behavior were evaluated using Pearson’s *r* correlation score. The effect size in the non-parametric statistics was measured as *r* = *z* / √*n*. Structural analyses were corrected for multiple comparisons using the false discovery rate (FDR) procedure (Benjamini and Yekutieli, 2001). The analysis approach was to use the fcMRI data as a discovery sample. ROIs identified with this approach were used to compute an fcMRI contrast map between A350V and WT, corrected for multiple comparisons using cluster-level correction. Only a single ROI and a single region survived this correction and it was subsequently correlated with behavior. Intergroup differences of the behavioral data were quantified using two-way repeated-measures ANOVA, with side preference (i.e. time spend in close interaction with *stranger 1*/object) defined as a repeated measure factor and genotype as between subject factor. Detection of outliers in the data was conducted following the steps suggested in Seo (2006). Symmetry and normality assumptions were assessed for selecting the appropriate detection method. Hence, Tukey’s and MAD (median absolute deviation) methods were used to detect and exclude outliers from the behavioral data.

### Data Sharing

All imaging raw data and the relevant codes used in this study will be made available in BIDS format on OpenNeuro, https://openneuro.org/datasets/ds#####, upon publication.

## Results

### Brain morphology alterations in A350V mouse model

Anatomical alterations have been reported in both humans with ASD and mouse models of ASD (Ecker et al., 2015; Ellegood et al., 2015). Therefore, we sought to examine whether there are anatomical volumetric differences in the brains of A350V mice relative to WT littermates by using manual segmentation and automatic non-linear registration of structural MRI. Manual quantification of volumes (*n* = 13 per group) showed a statistically significant increase in whole-brain volume in A350V mice as compared to WT littermates (**Fig. 2A;** Mann–Whitney U-test, U = 41, *p* = 0.027, *r* = 0.432), but no difference in ventricle volume (WT: 9.937 ± 0.418 mm^3^ [mean ± SEM]; A350V: 9.685 ± 0.281 mm^3^; U = 78, *p* = 0.758). After subtraction of the ventricle volume, the intracranial volume of the A350V mice (359.912 ± 3.405 mm^3^) was found to be significantly larger relative to WT littermates (348.918 ± 2.484 mm^3^; U = 41, *p* = 0.027, *r* = 0.432), suggesting that the increased whole-brain volume is unlikely to be due to hydrocephalus. Next, we used a deformation-based morphometry method (Andersson et al., 2007) to automatically evaluate changes in the volumes of the entire brain and substructures. Consistent with the manual quantification, whole-brain volume as expressed by the Jacobian deformation was significantly larger in A350V compared to WT (**Fig. 2A**; U = 33, *p* = 0.009, *r* = 0.512). In addition, a significant correlation between the manual whole-brain volume measurements and the Jacobian deformation measure (**Fig. 2B**; *n* = 13, *r*(24) = 0.845, *p* < 0.001) suggested that the automatic deformation measure with the current structural imaging acquisition parameters was reliable. The AMBC atlas was used to define the substructures of the brain and compare volumes of A350V to those of WT (**Fig. 2C**). We found increased volumes in A350V relative to WT in the cerebral cortex without the olfactory bulb (U = 22, *p* = 0.007; FDR-corrected, *r* = 0.623) and the hippocampal region (U = 31, *p* = 0.016; FDR-corrected, *r* = 0.533), but no significant volume differences in the thalamus, caudoputamen and piriform cortex (U values > 42, *p* values > 0.052; FDR-corrected).

**Figure 2.**
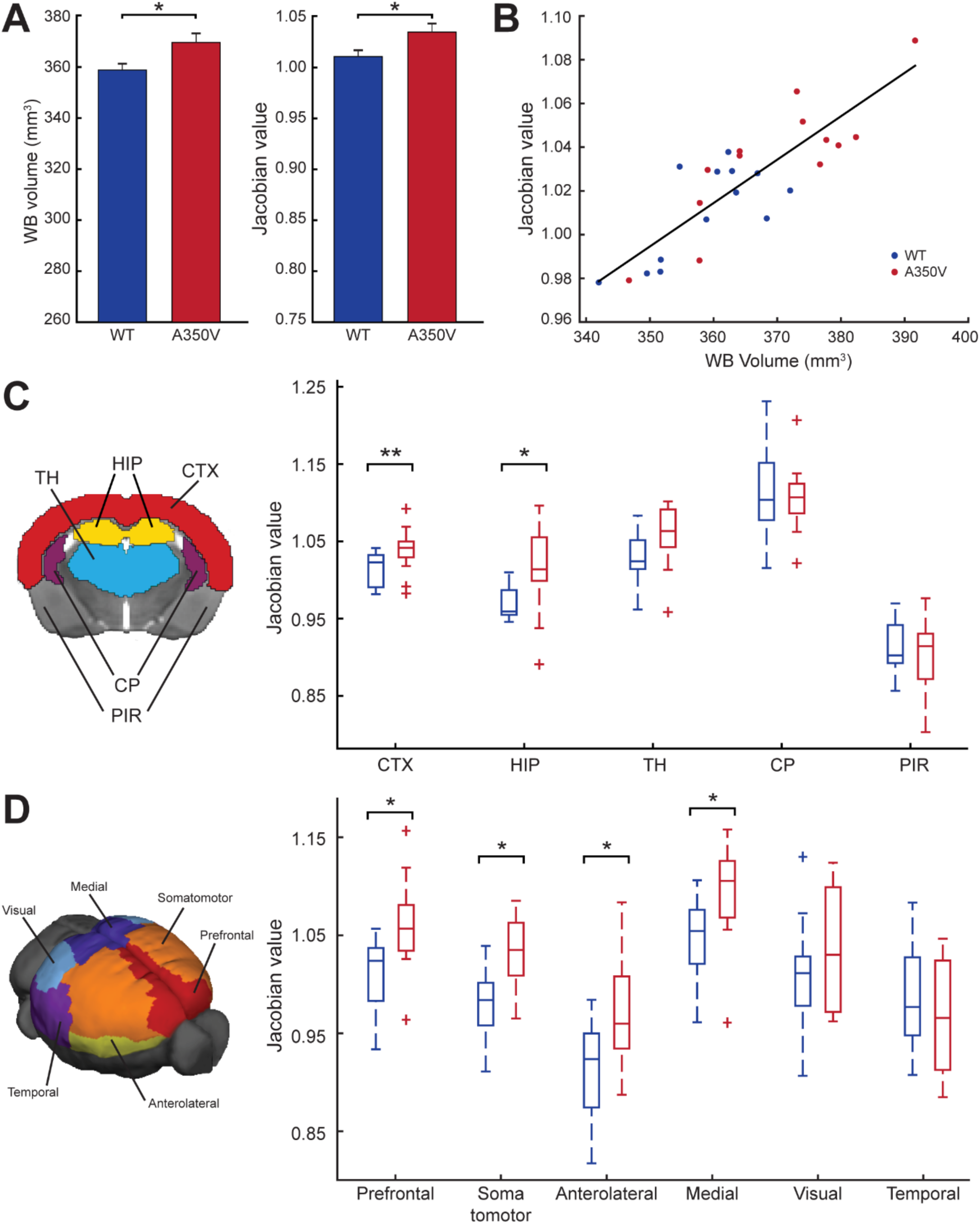
Structural alterations in A350V IQSEC2 mouse model. ***A***, Whole-brain volume increase in A350V mice shown by manual volume quantification (*left*) and automatic registration evaluation (*right*). ***B***, Correlation between Jacobian determinant and whole-brain volume showing an agreement between the two evaluation methods. ***C***, Jacobian determinant of the different brain structures depicted on the left indicates increased volume of CTX and HIP in A350V mice. ***D***, Jacobian determinant of the six cortical modules depicted on the left indicates increased volume of the Prefrontal, Somatomotor, Anterolateral and Medial modules in A350V mice. Data indicate mean ± SEM. Mann–Whitney U Test; * *p* < 0.05, ** *p* < 0.01, FDR-corrected. CTX, cerebral cortex; HIP, hippocampal region; TH, thalamus; CP, caudoputamen; PIR, piriform area.

Next, we tested whether the observed structural alterations are attributed to specific regions of the cortex by dividing it into six modules (Prefrontal, Anterolateral, Somatomotor, Visual, Medial and Temporal; **Fig. 2D**) following Harris et al. (2018). Volumes were significantly larger in Prefrontal (U = 30, *p* = 0.017; FDR-corrected, *r* = 0.543), Somatomotor (U = 30, *p* < 0.017; FDR-corrected, *r* = 0.543), Anterolateral (U = 39, *p* = 0.031; FDR-corrected, *r* = 0.452) and Medial (U = 35, *p* = 0.024; FDR-corrected, *r* = 0.492) modules in the A350V compared to WT (*n* = 13 per group), with no significant differences observed in the Visual (U = 67, *p* = 0.383) and Temporal (U = 69, *p* = 0.441) modules.

### Structure–function analysis reveals cortical alterations in A350V mice

To assess the functional disruption in the A350V mouse brain, we used functional connectivity MRI. It has been previously shown that fcMRI tracks anatomical connectivity, and that this coupling can be used to reveal functional connectivity based on brain architecture (Stafford et al., 2014; Bergmann et al., 2016; Grandjean et al., 2017; Melozzi et al., 2019). We used ROC analysis to evaluate the structure– function relations of cortical systems, by comparing the optical density tracer injection maps of the AMBC to the functional connectivity maps acquired using fcMRI (**Fig. 3A**). First, we examined the agreement between anatomical and functional connectivity at the group level, for systems that are hierarchically organized (sensory networks) relative to systems that are not predominantly hierarchical (association networks), as previously shown by Bergmann et al. (2016). This comparison is informative as it allows an assessment as to whether structure–function relations at the whole-brain level can be better explained by first synapse projections of sensory networks, due to their hierarchical organization, relative to association networks. At the group level, seed-based statistical parametric maps were computed for A350V and WT separately and then submitted to ROC analysis. We found that the area under the ROC curve (auROC), a measure of predictive strength, was higher in sensory relative to association networks for both groups (**Fig. 3B**; WT: U = 84, *p* = 0.022, *r* = 0.374; A350V: U = 71, *p* = 0.007, *r* = 0.442). There were no significant differences when comparing the auROC of each class of networks separately between A350V and WT (U_Sensory_ = 264, *p* = 0.628; U_Association_ = 75, *p* = 0.644). Finally, we sought to test whether the observed anatomical macrocephaly we found in A350V mice would be reflected in the structure–function relations of the six cortical modules. When comparing the auROC values of each module between the groups we found no significant differences (**Fig. 3C**; U_Prefrontal_ = 262, *p* = 0.965; U_Somatomotor_ = 413, *p* = 0.345; U_Anterolateral_ = 16, *p* = 0.8182; U_Medial_ = 37, *p* = 0.344; U_Visual_ = 230, *p* = 0.787; and U_Temporal_ = 35, *p* = 0.666).

**Figure 3.**
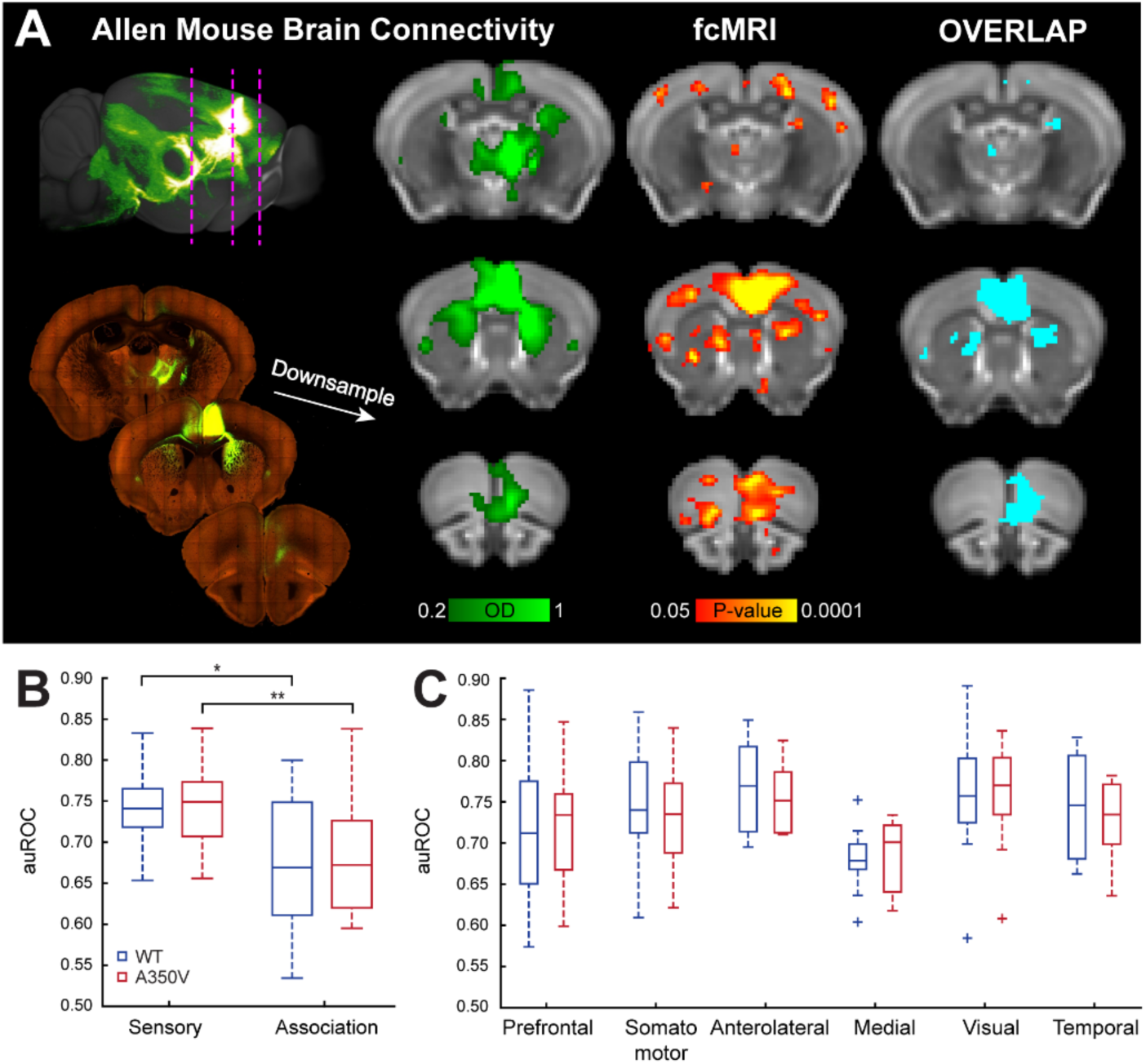
Anatomical connections predict functional connectivity at a group-level. ***A***, Analysis scheme for comparing structure–function relations using MRI shown for Allen mouse brain connectivity (AMBC) atlas experiment #112514202, ACAv injection. A maximum intensity projection sagittal view of signal density (*top, left*) and three representative coronal slices showing multiple anatomical connections (*bottom, left*) in AMBC space. Approximate locations of the presented coronal sections are indicated by magenta dashed lines. The arrow indicates the corresponding downsampled coronal slices (optical density, [OD] in green-light green, normalized OD > 0.2). An fcMRI statistical parametric map (in *red*-*yellow*) of positive correlations of ACAv on a downsampled version of the AMBC Atlas that matches the original fMRI resolution; *p* < 0.05, corrected for multiple comparisons using family-wise error rate correction for the whole mouse brain. Overlap of functional and anatomical connectivity (*pale blue*) demonstrates a close agreement across the two modalities. ***B***, The area under receiver operating characteristic (auROC) curve values demonstrate higher structure–function relations in sensory relative to association systems for both WT (*blue*) and A350V (*red*) mice. ***C***, The six cortical modules auROC values demonstrate equivalent structure–function relations between the two groups for all modules.

Quantification of structure–function relations allows an assessment as to whether disruption in functional connectivity at the single animal level localizes to subnetworks within the modules as defined by specific tracer injections and their first synapse projections. We adapted the above structure–function group-level approach to the individual level by submitting correlation maps (*Z*(*r*)) of each individual animal (*n* =13 per group), rather than the group average seed-based statistical parametric map (as was done above), to the ROC analysis. We first replicated the sensory vs. association networks analysis described above at the individual level by averaging all system-related seeds per animal. Similar to the group-level auROC analysis, we found that the sensory system was higher than the association system in both groups (**WT**: Sensory = 0.743 [median], Association = 0.676; Wilcoxon Signed-rank test, *p* < 0.001, Z = 3.18, *r* = 0.623; **A350V**: Sensory = 0.74, Association = 0.673; Wilcoxon Signed-rank test, *p* < 0. 001, Z = 3.18, *r* = 0.623).

Next, to identify specific disrupted regions we quantified structure–function relations for individual injections reported by Harris et al. (2018). We generated seeds that correspond to the locations of 101 C57BL/6J tracer injections, computed the seed-based correlation maps of individual animals and compared the auROC values of each injection across the two groups. We discovered significant differences in six injections located in the anterior cingulate, auditory and visual cortices (**Fig. 4**). The auROC values for the ACAd and ACAv regions were significantly higher in A350V relative to WT (U_ACAd_ = 42, *p* = 0.031, *r* = 0.422; U_ACAv_ = 34, *p* = 0.01, *r* = 0.503), suggesting that the anatomical connectivity served as a better predictor than the functional connectivity in A350V. Conversely, AUDpo, AUDd, VISli and VISp regions showed reduced auROC values in A350V relative to WT (U_AUDpo_ = 42, *p* = 0.031, *r* = 0.422; U_AUDd_ = 24, *p* = 0.002, *r* = 0.603; U_VISli_ = 42, *p* = 0.031, *r* = 0.422; U_VISp_ = 42, *p* = 0.031, *r* = 0.422).

**Figure 4.**
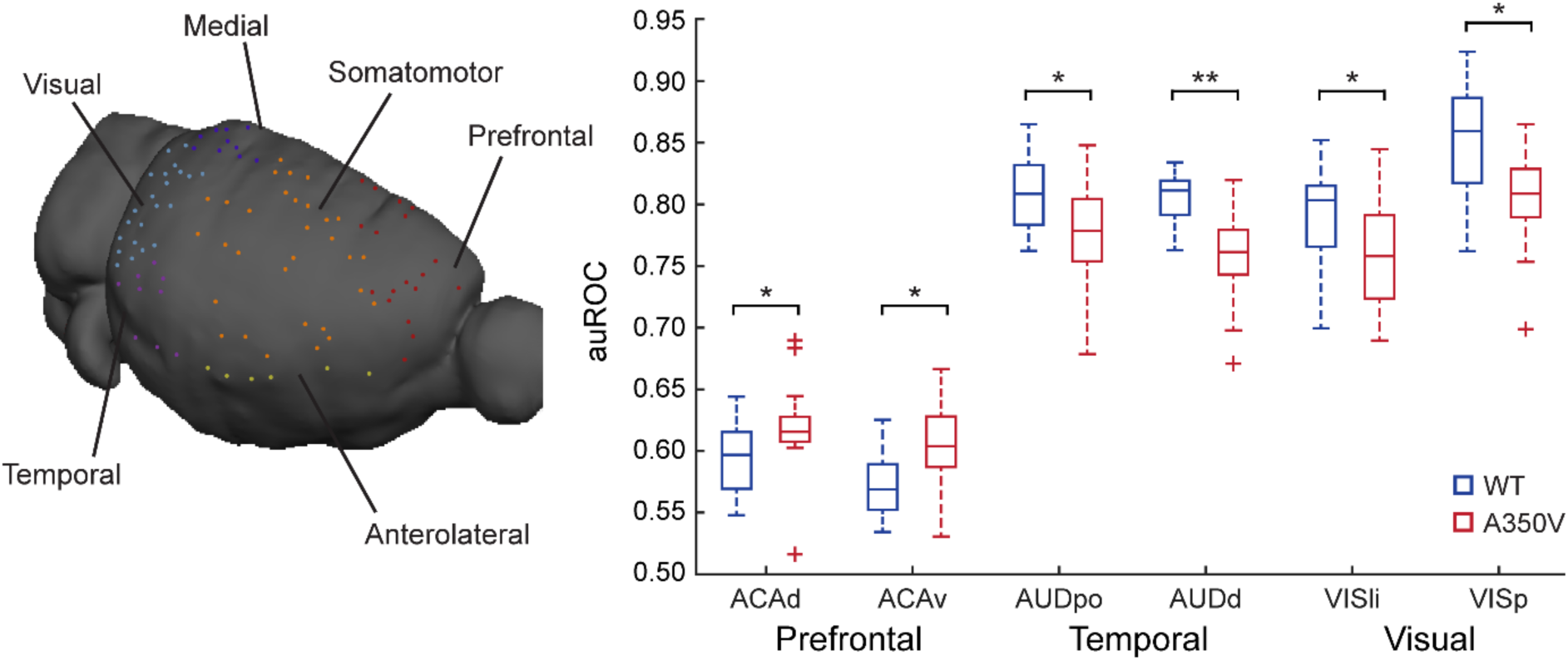
Structure–function relations at the individual level reveal altered functional connectivity in A350V relative to WT. The seeds corresponding to AMBC tracer injections are illustrated on a mouse cortical surface (*left*). Seeds are color coded for the six cortical modules. The area under receiver operating characteristic (auROC) curve values (*right*) indicate increased structure–function relations in A350V mice (*red*) for ACAd and ACAv and greater overlap in WT mice (*blue*) for AUDpo, AUDd, VISli and VISp. Mann–Whitney U Test; *n* = 13 per group, * *p* < 0.05, ** *p* < 0.01 (uncorrected for multiple comparisons). ACAd, anterior cingulate area, dorsal part; ACAv, anterior cingulate area, ventral part; AUDpo, posterior auditory area; AUDd, dorsal auditory area; VISli, laterointermediate visual area; VISp, primary visual area.

Following the altered structure–function relations described above, we sought to further investigate whether these regions demonstrated functional changes at the network level. Since the pairs of the regions (ACAd/ACAv, AUDpo/AUDd, and VISli/VISp) are potentially overlapping to the extent that they are effectively identical, we evaluated the overlap of the tracer injections for each pair of proximal regions using the Sørensen–Dice similarity coefficient. We found a substantial overlap for ACAd/ACAv and AUDpo/AUDd (Dice index of 0.861 and 0.757, respectively), and therefore in further analyses they were considered as single regions, designated ACAd/v and AUDpo/d. The VISli and VISp similarity coefficient was 0.545 and they were therefore considered as two separate networks.

### Corticostriatal alterations in functional connectivity are associated with social impairments

To further dissect the alterations found in the structure-function analysis we computed a two-way *t*-test statistical parametric map between the groups for each of the above seed-based correlation maps. Of the above regions, only the comparison between A350V and WT of the ACAd/v seed-based maps showed a significant increase in functional connectivity in A350V mice which was exclusive to the dorsomedial caudoputamen (CP) (**Fig. 5A;** *t*-test, *p* = 0.01, FWE cluster-level corrected, with cluster-defining threshold of *t*(24) > 2.49, *p* < 0.01). Next, we performed a seed-to-seed correlation analysis between the ACAd/v and the ROI in the CP that was found in the prior analysis obtaining correlation values at the individual animal level (**Fig. 5B**). As expected, A350V mice displayed increased ACAd/v– CP connectivity relative to WT (U = 26, *p* = 0.002, *r* = 0.583).

**Figure 5.**
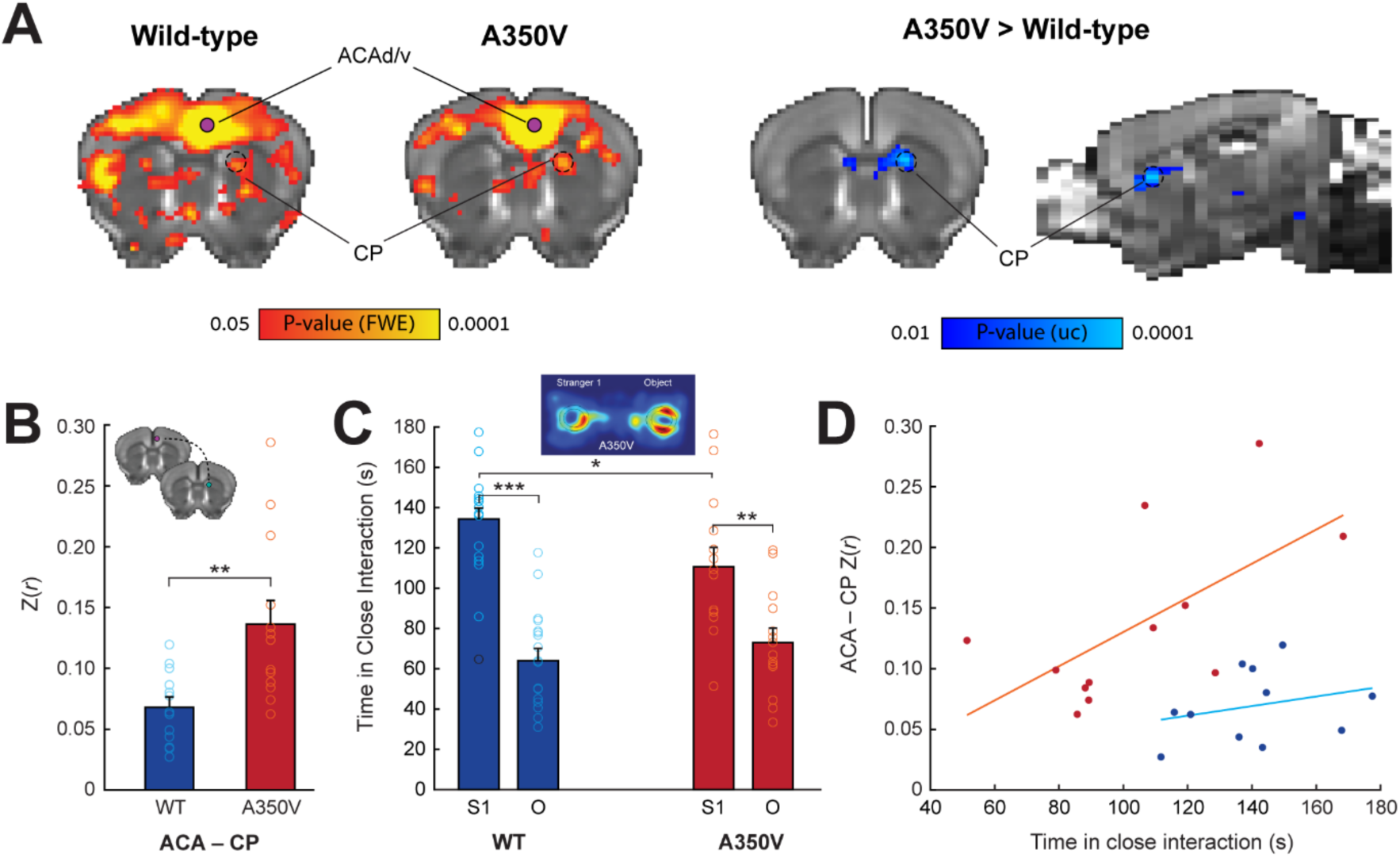
Corticostriatal functional connectivity correlates with disrupted sociability in A350V mice. ***A***, Seed-based comparison of the ACAv/d showing fcMRI statistical parametric maps (*red-yellow*) of positive correlations for WT (*left*) and A350V mice (*right*) (*p* < 0.05, corrected for multiple comparisons using whole-brain family-wise error (FWE) rate correction) and an fcMRI statistical parametric map (*blue-light blue*) of increased connectivity in A350V relative to WT showing significant increase in the dorsomedial CP (two-way unpaired *t*-test, *p* = 0.01, FWE cluster-corrected, with cluster-defining threshold of *t*(24) > 2.49, *p* < 0.01, voxel extent ≥ 5). All maps are displayed on a downsampled version of the AMBC Atlas that matches the original fMRI resolution. Purple circles indicate the ACAd/v seed location. Black dashed circle indicates CP cluster that was found significant. ***B***, Seed-to-seed correlation between ACAd/v and CP showing increased connectivity in A350V mice. Colored circles indicate individual subject z-score. Coronal slices showing seed location for ACAd/v (*purple*) and CP (*green*) (*top left*). ***C***, Duration of time spent in close interaction with Stranger 1 (S1) and an object (O) in the three-chamber social preference test demonstrating that both A350V and WT prefer interacting with S1, although this preference is less pronounced in A350V. Colored circles indicate individual subject variability in sociability measurement with black circle indicating outlier in the data. A representative heat map analysis of A350V mouse is depicted (*top center*). ***D***, Connectivity between ACAd/v and CP in relation to sociability showing significant correlation for A350V mice only. Data indicate mean ± SEM. Mann–Whitney U Test (**A**); ** *p* < 0.01. Two-way rmANOVA, with Holm-Bonferroni post hoc t-test (**C**); * *p* < 0.05, ** *p* < 0.01, *** *p* < 0.001. ACA, anterior cingulate area; CP, caudoputamen.

In the human index case with the A350V IQSEC2 mutation abnormalities in social behaviors were found (Zipper et al., 2017). Thus, social behavior was assessed using a three-chamber sociability test (**Fig. 5C**). When we examined social preference as a measure of time spent in close interaction, a repeated-measures ANOVA revealed a significant main effect of Side (F(1,29) = 45.39, *p* < 0.001, η _p_ ^2^ = 0.61) as well as an interaction between Side and Genotype (F(1,29) = 4.17, *p* = 0.05, η _p_ ^2^ = 0.126). A further post-hoc analysis revealed a significant difference in time spent in close interaction with an unfamiliar mouse between A350V and WT (*t*-test, *t* = 2.444, *p* = 0.035, Holm–Bonferroni correction). Collectively, these results indicate that both A350V and WT mice prefer interacting with an unfamiliar mouse over an object yet this preference was reduced in A350V relative to WT.

Finally, to test whether the abnormal social behavior found in A350V model can be explained by the neurophysiological alterations that were obtained from the same animal cohort, we correlated the social preference measurement both with the structural deformation and the seed-to-seed functional connectivity. The observed volumetric differences showed no significant correlation with the behavioral impairments that were found (data not shown). When examining the relations between the altered ACAd/v–CP functional connectivity with the observed social behavior, we found a significant correlation of functional connectivity magnitude and social preference for A350V only (**Fig. 5D;** A350V: *r*(10) = 0.627, *p* = 0.029; WT: *r*(9) = 0.267, *p* = 0.426). Namely, equivalent functional connectivity between A350V and WT resulted in reduced social interaction in A350V, while increased corticostriatal connectivity in A350V is associated with normal levels of social interaction as found in WT.

## Discussion

We explored the neurophysiological changes in a novel transgenic A350V mouse model which is based on a human *de novo* mutation in the IQSEC2 gene associated with ASD. We show that A350V mice exhibit overall increased brain volume, restricted to the cortex including the hippocampus, along with alterations in structure–function relations of the frontal, auditory and visual systems. We also show changes in corticostriatal functional connectivity which are associated with individual variability in social behavior, demonstrating that the equivalent levels of corticostriatal functional connectivity between A350V and WT correspond to reduced social interaction in the three–chamber social preference test in A350V. Collectively, these findings suggest that the A350V missense mutation in the IQSEC2 gene induces ASD features that are derived from corticostriatal dysfunction and may serve as a translational model from human to mouse.

IQSEC2 is one of many ASD-linked genes that encode post synaptic density proteins of glutamatergic synapses, such as CNTNAP (contactin-associated protein), neuroligin, NF1 (neurofibromin 1), SHANK family and more (Peça and Feng, 2012). While the signal transduction pathway of IQSEC2 is not fully understood, it has been previously suggested that IQSEC2 functions as a guanine nucleotide exchange factor (GEF) for ADP-ribosylation factors (Arfs), and that the Arf-GEF activity further regulates AMPA receptor trafficking (Murphy et al., 2006; Brown et al., 2016). In the case of the A350V mutation, it has been proposed that there is a constitutive activation of the GEF activity of IQSEC2 that results in increased endocytosis of AMPA receptors via constant activation of Arf6 along with other processes that are less understood (Rogers et al., 2019).

In ASD, human studies have consistently demonstrated a modest but significant increase in overall brain volume in toddlers (Piven et al., 2017). In a recent study characterizing structural alterations in 26 ASD mouse models using MRI, these models were classified to three subtypes according to alterations in volume and connectivity (Ellegood et al., 2015). The authors found that all of the mouse models that were associated with increased brain volume were classified in the same subtype (NRXN1α^-/-^NRXN1α^-/+^ andFMR1-/yFVB). However, other models within this subtype either showed no change in brain volume (En2^-/-^ and FMR1^-/y^ B^) or decrease in brain volume (SHANK3^-/-^ and SHANK3^-/+^). Collectively, this evidence suggests that brain volume is not a critical distinguishing feature. Related to the reported A350V mutation, in a recent study using female mice (Jackson et al., 2019), heterozygous loss of function of the IQSEC2 gene showed thinning of the corpus callosum along with increased volume of the hippocampus and specifically the dentate gyrus within it (cortical volume was not reported). Here, male A350V mice presented an increase in whole-brain volume that was found to derive from cortical and hippocampal enlargement. We found a volume increase in four out of the six cortical modules, suggesting a non-specific cortical enlargement. Further, unlike the functional connectivity results, increased volume was not correlated with altered social behavior identified in this model and are possibly relevant to other behavioral features presented in this model (Rogers et al., 2019). Taken together with Ellegood et al. (2015), it is suggested that the observed macrocephaly in A350V mice is unlikely to be involved in the pathophysiology of the brain connectivity we report here. Numerous studies implicated corticostriatal alterations in ASD pathophysiology in humans and animals. A study comparing ASD and typically developing children showed increased functional connectivity between striatal subregions and both association and limbic cortices (Di Martino et al., 2011). In a study using connectivity-based parcellation, a difference in the organization of corticostriatal circuitry in ASD was reported, demonstrating that the separation of the putamen to distinct anterior and posterior clusters is absent in ASD participants, and is potentially driven from a connectivity fingerprint of several cortical regions (Balsters et al., 2018). Previous studies in the SHANK3 mouse model have demonstrated dysfunction of corticostriatal activity (Peça et al., 2011; Peixoto et al., 2016), with the striatum playing a causal role in repetitive grooming behavior (Wang et al., 2017). A recent fcMRI study in the same mouse model showed an overall prefrontal hypoconnectivity along with disrupted functional connectivity between the ACA and the caudoputamen (Pagani et al., 2019). Consistent with these reports, we found altered functional connectivity between the ACA and the dorsomedial striatum. Collectively, these findings emphasize the contribution of association and limbic corticostriatal hyper- or hypo-connectivity to ASD-related pathology. Moreover, anatomical homology and recapitulation of findings when using fMRI in ASD genetic mouse models suggest that the pathophysiological findings in genetic animal models are likely to translate to humans.

Studying the contribution of specific brain structures to behavior is particularly important when it comes to understanding neuropathology of diseases, as it can help dissect the relevance of specific brain structures to behavioral features and subsequently identify disease pathophysiology. In a task-based fMRI study, hyperconnectivity between the anterior cingulate cortex and the caudate has been shown to be associated with deactivation to social rewards in ASD participants (Delmonte et al., 2012). Several studies of ASD mouse models linked brain function and social behavior impairments. Homozygous loss of the *CNTNAP2* (contactin-associated protein-like 2) gene results in frontoposterior hypoconnectivity, a finding that has been reported in several ASD models, which was further associated with reduced social investigation in a male–female social interaction test (Liska et al., 2018). In addition to ACA–CP disrupted connectivity, the SHANK3 fcMRI study mentioned above showed that prefrontal hypoconnectivity is correlated with socio-communicative deficits (Pagani et al., 2019). These findings are consistent with a study showing that specific deletion of SHANK3 in the ACA is sufficient to induce social impairments, suggesting that the ACA plays a role in regulating social behavior (Guo et al., 2019). In our study, A350V mice presented high variability in the sociability measurement using the three-chamber sociability task. We found this variability to be correlated with the connectivity levels between the ACA and the dorsomedial striatum. While the A350V mouse model showed an overall increase in corticostriatal connectivity, we observed that the A350V mice with functional connectivity levels equivalent to WT littermates presented low sociability levels as compared to these WT littermates, while the A350V mice with high connectivity levels presented normal sociability levels. This result suggests that increased levels of connectivity may reflect a compensatory neural response mitigating social impairments in the A350V model. The concept of neural response of compensatory nature on cognitive function has long been recognized (Basso et al., 1989; Ullman and Pullman, 2015) and has been previously suggested to underly variability in disease severity in ASD (e.g., Livingston and Happé, 2017). To fully understand the pathophysiology of the A350V model future studies will need to causally address the role of ACA in social impairments.

Finally, we would like to propose the A350V IQSEC2 mouse as a putative ASD model. Overall, our results are consistent with brain alterations found in several ASD models. The diversity of genetic mutations which lead to similar brain-wide and associated behavioral changes in ASD suggests that the ability to offer precise therapies for ASD depends on understanding how diverse neural pathologies are linked to ASD. Specifically, future research which will more fully characterize the cellular pathophysiological changes that drive the brain-wide level effects observed in the A350V model and the relation of these changes to the brain-wide changes observed across the diverse ASD models will facilitate progress in understanding the causes of ASD in A350V IQSEC and proposing paths to therapy.

## Acknowledgments

This work was supported by the Israel Science Foundation (770/17; to I.K.), the National Institutes of Health (1R01NS091037; to I.K.) and the Adelis Foundation (to I.K.). We thank Technion’s Biological Core Facilities and Edith Suss-Toby for her assistance with MRI, and the Technion Preclinical Research Authority for assistance with animal care.

